# The Role of Video Game Practice in Trial-by-Trial Adaptation, Consolidation, and Reinforcement Learning Biases

**DOI:** 10.1101/2025.08.24.672020

**Authors:** Luis A. Llamas-Alonso, Andrea J. Quevedo-Calderon, Sandra L. Quiñones-Beltran, Ana Lucía Jiménez-Pérez, Arturo Arvizu-Oviedo, Armando Q. Angulo-Chavira

**Affiliations:** Facultad de Ciencias Administrativas y Sociales, Universidad Autónoma de Baja California, Ensenada, México; Facultad de Psicología, Universidad Nacional Autónoma de México, Ciudad de México, México

**Keywords:** Video games, Reinforcement learning, Feedback adaptation, Probabilistic Selection Task, Learning Dynamics

## Abstract

This study examined whether habitual video game play influences reinforcement learning dynamics, feedback adaptation, consolidation, and motivational biases. Two groups of participants (gamers and controls) completed a Probabilistic Selection Task assessing learning from positive and negative feedback across three phases: learning, test, and transfer. Mixed-effects modeling revealed that gamers showed enhanced learning trajectories, particularly under high-uncertainty conditions, greater sensitivity to accumulated rewards, and a more exploitative decision pattern compared to controls. In the test phase, gamers demonstrated higher accuracy, especially on difficult stimulus pairs, suggesting superior consolidation of probabilistic value representations. However, no group differences emerged in transfer-phase approach/avoidance biases or decision consistency. These findings suggest that habitual video game experience enhances dynamic feedback integration and value updating in uncertain contexts, but does not alter stable reinforcement learning biases. Results have implications for leveraging game-like environments to enhance adaptive learning strategies in educational and clinical settings.

## Introduction

Video games offer a rich environment for adaptive learning. Like many real-life situations, they require making decisions under uncertainty and adjusting behavior based on ambiguous or delayed feedback——much like choosing your commute route with incomplete traffic information and then updating the next day based on whether today’s choice saved time (reward) or caused a delay (penalty). These dynamics resemble the structure of reinforcement learning tasks used in cognitive science, where behavior is shaped by the consequences of past actions [1]. Reinforcement-based learning involves tracking the value of choices across trials, allowing individuals to optimize future outcomes by learning from both rewards and penalties [2,3]. Given these parallels, video games can be regarded as ecologically valid contexts that instantiate core reinforcement contingencies; thus, studying individuals with video game experience can provide insights into how these environments shape reinforcement learning.

Although the principles of reinforcement learning are universal, individuals differ in their sensitivity to different types of consequences. Some people are more responsive to rewards, biasing learning toward approach; others are more sensitive to punishment, biasing learning toward avoidance. These motivational learning tendencies have been linked to stable traits such as reward sensitivity and punishment sensitivity, which shape how people interpret and react to environmental feedback [4,5].

To study how individuals develop preferences for learning from positive or negative feedback, researchers have frequently employed the Probabilistic Selection Task (PST;[2]). In this task, participants learn to discriminate between abstract symbols based on probabilistic reinforcement schedules during a training phase, followed by test and transfer phases where choices are made without feedback. The transfer phase is commonly interpreted as reflecting stable learning biases, specifically a tendency to choose the most positively reinforced option (approach learning) or to avoid the most negatively reinforced one (avoidance learning). This paradigm has proven useful for capturing individual variation in feedback processing and has been applied across a range of cognitive and clinical studies [3,6].

Yet, while the transfer phase provides a snapshot of consolidated preferences, focusing solely on final performance may obscure the dynamic learning processes that precede it. Trial-by-trial behavior during earlier training and test phases captures how participants integrate feedback, adjust expectations, and adapt decision strategies over time. This incremental adaptation lies at the core of reinforcement learning theory, where prediction errors (the discrepancy between expected and actual outcomes) guide the continuous updating of value representations [1]. Understanding how individuals modify their choices in response to ongoing reinforcement may reveal important aspects of learning flexibility, early sensitivity to feedback, and the stabilization or reversal of strategies [7]. Thus, a more comprehensive view of motivational learning biases must consider the temporal trajectory of behavior across all phases of the task, not just its endpoint.

In this regard, the Weather Prediction Task (WPT) examines probabilistic categorization learning, in which participants gradually predict outcomes from probabilistic cue configurations [8]. It has been useful for probing how declarative and feedback-based systems support performance under uncertainty, and Schenk and colleagues reported that video-game players outperform non-players on this task. However, the WPT does not assess consolidated performance when feedback is withheld, nor does it dissociate learning driven by rewards versus punishments.

By contrast, the Probabilistic Selection Task (PST) provides a complementary framework for reinforcement learning: it supports trial-by-trial modeling of value updating during acquisition, tests consolidation when feedback is withheld, and quantifies transfer-phase biases (e.g., approach/avoid tendencies such as “choose-A” vs. “avoid-B”; [2]). Because commercial games rely on repeated feedback and behavioral adaptation, the PST is well suited to ask not only whether gamers learn more efficiently, but how—for example, via enhanced reward sensitivity, improved feedback tracking, or a stronger approach-oriented decision policy— mechanisms that the WPT cannot isolate.

Importantly, the way reinforcement contingencies are constructed within the PST also shapes learning dynamics. Most studies focus on high-contrast stimulus pairs (e.g., 80/20 reward probability), assuming these provide the clearest signal for assessing learning [2,3,6]. Yet, this emphasis may oversimplify the complexity of real-world learning, where feedback is often ambiguous or inconsistent. Discriminating between low-contrast pairs (e.g., 70/30 or 60/40) requires greater sensitivity to probabilistic structure and may engage more effortful cognitive processing, making these conditions valuable for detecting differences in feedback integration, decision noise tolerance, and reinforcement sensitivity [2,3,9].

Prior work supports this possibility. Video game players have been shown to exhibit heightened responsiveness to reward-related cues and faster adaptation to reinforcement contingencies [10]. Moreover, video gameplay engages the mesolimbic-cortical dopaminergic system, central to reward processing and reinforcement learning [11,12].

Repeated activation of this circuitry may induce neuroplastic changes that amplify reward sensitivity and bias learning toward approach-based strategies. These mechanisms, if validated, would illuminate how motivational learning tendencies emerge and adapt in interactive, feedback-rich environments.

Understanding these dynamics is especially relevant given the role of motivational learning biases in both clinical and educational outcomes. Heightened punishment sensitivity has been linked to anxiety and compulsive behaviors, while excessive reward orientation may underlie impulsivity and self-regulation difficulties [13,14]. In academic contexts, students’ motivational dispositions—such as their perceived value of feedback [15] and their ability to delay gratification and maintain a future time perspective [16] can shape how they respond to corrective input. Those with low motivation toward feedback or a limited orientation toward future goals may be less inclined to engage with feedback that requires sustained effort before yielding benefits, resembling avoidance-oriented patterns. In contrast, students who value feedback highly and respond readily to immediate input may benefit when reinforcement is rapid but find it more challenging to sustain engagement when rewards are delayed. These findings underscore the importance of identifying contextual factors that modulate sensitivity to reward and punishment during learning. One such factor may be prior exposure to environments like video games, which, due to their reliance on feedback-driven challenges, variable reinforcement schedules, and goal-directed action, could influence how individuals acquire and consolidate learning over time.

The present study aimed to examine whether reinforcement learning dynamics—specifically, trial-by-trial adaptation to positive versus negative feedback, followed by consolidation and transfer without feedback—are modulated by habitual video game use. We hypothesized that frequent gamers would exhibit greater sensitivity to reward-related cues and enhanced behavioral flexibility during the learning phase, reflected in faster acquisition and more efficient adjustment to probabilistic feedback, particularly under ambiguous reinforcement conditions. In the test phase, we expected gamers to demonstrate superior discrimination performance, especially on low-contrast stimulus pairs, indicative of improved consolidation of probabilistic value information. Finally, in the transfer phase, we predicted that gamers would exhibit stronger approach-oriented learning biases over avoidance-oriented biases and greater choice consistency, reflecting more stable consolidation of reinforcement learning preferences. To test these predictions, we adapted the Probabilistic Selection Task into a game-like format and analyzed performance across all phases, with a specific focus on probabilistic contrasts that vary in feedback clarity.

## Methods and Materials

### Participants

The present study included a total of 72 young adults, comprising 36 non-players (Control) (17 males; aged *M*= 22.17, *SD* = 3.02; and 19 females aged *M* = 21.78, *SD* = 2.46) and 36 video game players (Gamer) (19 males; *M* = 21.84, *SD* = 2.50; and 17 females; *M*= 21.67, *SD* = 2.56). Primary game genres for the gamer group are presented in Table S1.

Participants were actively enrolled college students who completed a brief clinical history questionnaire; none reported medical, neurological, or psychological conditions that could compromise cognitive performance. Recruitment materials stated that final performance would be ranked, with the top three earning online gift cards worth $800, $500, and $300 MXN. This incentive was intended to enhance motivation and optimize task engagement.

Group classification was determined using a custom-designed questionnaire developed by the authors, currently undergoing validation, titled the Video Game Player Inventory, which assesses video game consumption habits. The Gamer group were required to actively play video games for seven or more hours per week maintaining that frequency for at least six months prior to the assessment. In contrast, Control group were required to play between zero and five hours of weekly gameplay and had no history of regular gaming at any prior life stage. All participants had normal or corrected-to-normal vision and reported no current or past psychological, psychiatric, or neurological conditions that could interfere with the study.

To characterize the sample and screen for variables that might influence the results, we measured: impulsivity level (Barratt Impulsiveness Scale -BIS-11-) and sensitivity to punishment and reward (The Sensitivity to Punishment and Sensitivity to Reward Questionnaire -SPSRQ-). All participants provided written informed consent. The research protocol was reviewed and approved by the Ethics Committee of the Universidad Autónoma de Baja California (ID: 430/2025-1).

### Stimuli and task

To assess reinforcement-based learning, the Probabilistic Selection Task proposed by Frank et al. (2004) was adapted and implemented using PsychoPy® version 2024.2.1, an open-source software for behavioral experiments [17]. This task evaluates whether individuals tend to learn better through positive feedback/reward (approach learning) or negative feedback/punishment (avoidance learning). To increase engagement, the probabilistic selection task was adapted into a game-like format in which participants were asked to decode symbols from an ancient language that we called “*The Codebreakeŕs Challenge*”. (The exact instructions given during every phase is provided in the S2). The task consists of three distinct phases: learning, test, and transfer, described below. The duration of the task ranged from 15 to 60 minutes, depending on the participant.

In the first learning phase participants were shown one of three symbol pairs (A/B, C/D, E/F) for 3500 ms. Each symbol was associated with a specific reinforcement probability (Figure 1). In the A/B pair, choosing symbol A was linked to an 80% chance of receiving positive feedback and a 20% chance of negative feedback; conversely, choosing symbol B had a 20% chance of positive feedback and an 80% chance of negative feedback. Similarly, the C/D and E/F pairs were associated with reinforcement contingencies of 70/30 and 60/40, respectively (Figure 1). After pressing the left or right key, the selected symbol was highlighted in blue on the screen for 300 ms, followed by positive or negative feedback displayed for 500 ms. Participants were shown their current score in the top-right corner, the number of remaining attempts in the top-left corner, and whether they gained or lost points in the center of the screen. A correct response earned ten points, while an incorrect one resulted in a ten-point deduction. Participants were instructed to use the feedback to learn which symbol to choose in order to maximize their score; however, they were not informed that each stimulus pair had associated reinforcement probabilities. If the response time exceeded 3500 ms, a message appeared on the screen prompting the participant to respond faster, and the trial was not scored. Each stimulus pair was presented 20 times in random order, totaling 60 trials [2,6].

**Figure 1.**
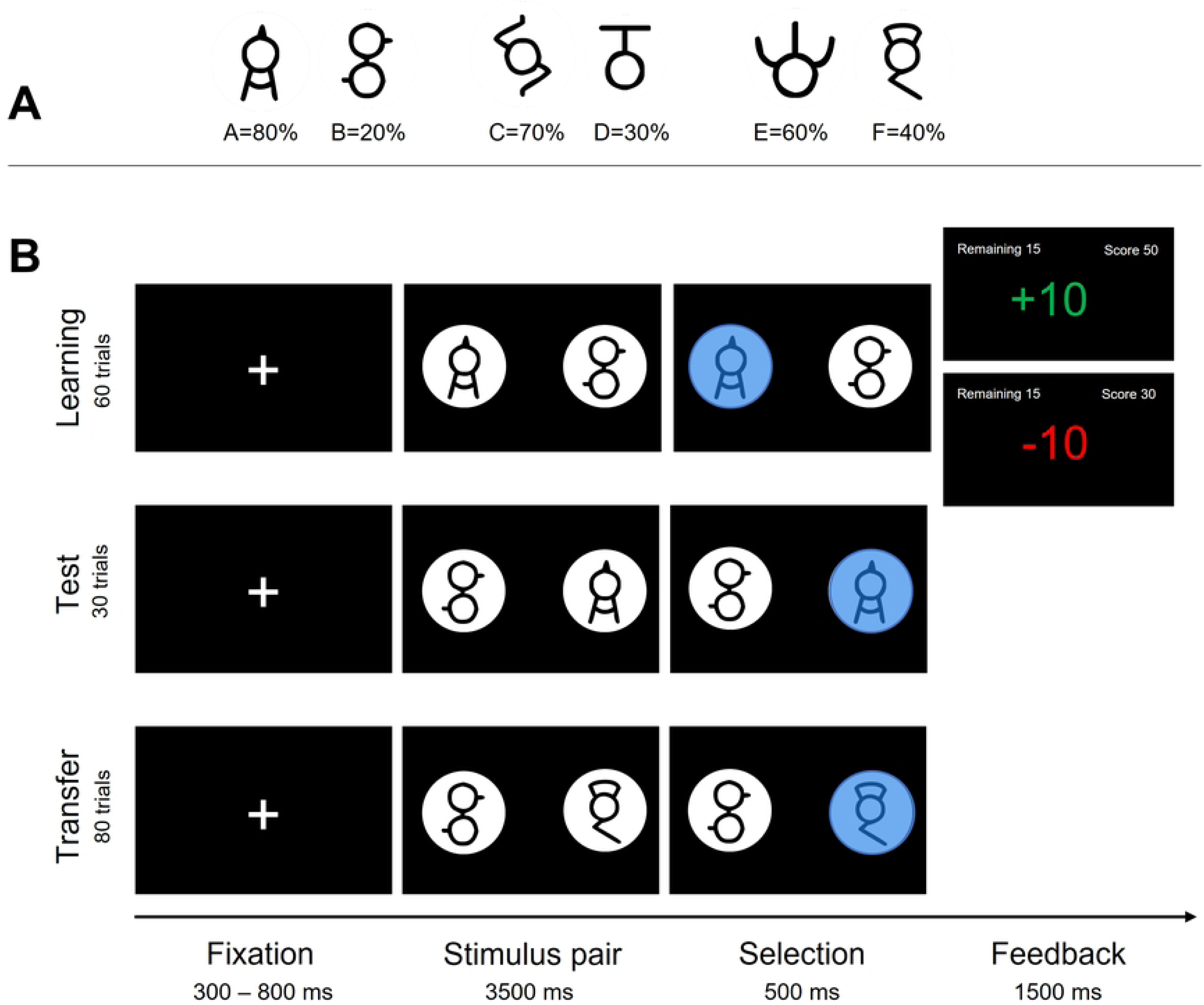
A) Symbol pairs and their associated reward probabilities. B) Example trial sequences for each phase of “The Codebreaker’s Challenge,” an adaptation of the Probabilistic Selection Task.

After completing the learning phase, participants proceeded to the test phase. The task structure remained the same, except that no feedback was provided, requiring participants to rely on the stimulus–outcome associations acquired earlier. The purpose of this phase was to assess whether participants continued to respond based on prior learning in the absence of feedback. Each stimulus pair was presented 10 times in random order, resulting in 30 trials. The transfer phase began once participants achieved at least 80% accuracy in the test phase—that is, 240 correct responses out of 300. If this criterion was not met, the learning and test phases were repeated after an optional one-minute break. Each participant could repeat these phases up to five times; if the accuracy threshold was still not reached, the transfer phase proceeded regardless [2,6].

In this final phase, the original stimulus pairs were dissolved and new combinations were formed. The symbol with the highest probability of positive feedback (“A”) was paired with all symbols it had not previously been paired with, resulting in the combinations: A/C, A/D, A/E, and A/F. Similarly, the symbol with the lowest probability of positive feedback (“B”) was paired with the remaining symbols to form the combinations: B/C, B/D, B/E, and B/F. The transfer phase consisted of 80 trials in total, with symbols A and B appearing in 40 trials each. No feedback was provided, and the stimulus sequence and timing were identical to those in the test phase [2,6].

### Data Processing and Statistical Analysis

All statistical analyses were conducted using R (version 4.3.1). Key packages included lme4 for fitting linear and generalized linear mixed-effects models. Mixed-effects models were optimized using the bobyqa optimizer when necessary. Statistical inference relied on Wald tests for fixed effects in mixed models and Welch’s t-tests for between-group comparisons where appropriate.

Participants with more than 20% missing trials were excluded from subsequent analyses. Reaction-time (RT) distributions were screened for within-subject outliers: RTs with absolute z-scores exceeding 2.5 were set to missing to prevent extreme values from biasing further analyses. No other trimming was applied.

### Learning Phase Analysis

First, we evaluated how many learning–test cycles participants completed before reaching the transfer criterion. The number of completed learning–test cycles was compared across groups via t-test, providing insight into how quickly each group met the learning criterion.

Performance across the Learning phase was analyzed by binning the single trials into blocks of ten trials. Mean accuracy per block was calculated separately for each participant and stimulus pair (A/B, C/D, E/F). A generalized linear mixed-effects model (GLMM) was then fitted to these accuracy data, including fixed effects for Group (Control vs. Gamer), Block (linear term), Pair (A/B, C/D, E/F), and all their interactions. Participant-specific random intercepts and random slopes for Block were included to accommodate variability in baseline performance and learning rates.

Additionally, we examined Win–Stay/Lose–Shift (WSLS) decision patterns. Each trial’s chosen symbol, outcome (win = +1, loss = -1, zero = 0), and stimulus pair context were recorded. A binary “stay” response was defined as repeating the previous choice within the same pair context. To account for the broader reward context beyond the immediate previous outcome, we computed an accumulated reward as the running average reward experienced by each participant up to the previous trial, standardized as a within-subject z-score. WSLS data were analyzed using a mixed-effects logistic regression, including fixed effects for Group, previous outcome (Win/Loss), cumulative GRS, and their interactions, alongside stimulus Pair. Random intercepts and slopes for the previous outcome were included by participant.

We also analyzed the exploration–exploitation trade-off throughout the Learning phase. Choices were categorized as exploitation if the participant selected the symbol with the higher programmed reward probability (A, C, E) or as exploration if the alternative lower-value symbol (B, D, F) was chosen. Trials without a recorded response or involving symbols outside the canonical pairs were discarded. Exploration probability was modeled using logistic mixed-effects regression, with trial number as a continuous predictor and group as a categorical factor. Random intercepts captured individual differences in baseline exploration rates.

### Test Phase Analysis

Given the absence of feedback during the Test phase, we directly assessed participants’ learned preferences by coding accuracy as choosing the higher-value symbol within each pair (A in A/B, C in C/D, E in E/F). Trials without a recorded response were omitted.

Accuracy data were analyzed with logistic mixed-effects regression, including fixed effects for Group, Pair, and their interaction, with participant-specific random intercepts.

To complement this, decisional consistency was evaluated using Shannon entropy of participants’ choices within each pair, quantifying the uncertainty or variability of their decisions. For each participant, we tallied symbol choices within each stimulus pair, converted these counts to probabilities, and calculated entropy:

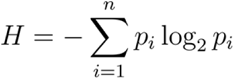

Entropy scores were analyzed using a linear mixed-effects model with Group, Pair, and their interaction as fixed effects, alongside random intercepts by participant.

### Transfer Phase Analysis

In the Transfer phase, previously learned pairs were recombined, and choices reflected internalized stimulus values. Analysis focused exclusively on trials involving stimuli A or B. For these trials, we coded “choose A” (selecting A when available) and “avoid B” (selecting any alternative when B was available). Each measure was analyzed using logistic mixed- effects regressions with Group as a fixed effect and random intercepts by participant.

To quantify motivational bias towards reward versus punishment, we computed the Valence Learning Bias Index (VLBI) for each participant as the difference between their proportion of A selections and B avoidances. Positive VLBI values indicate a bias toward reward-driven approach, while negative values indicate a bias toward punishment-driven avoidance. Group differences in VLBI were analyzed using a simple linear regression.

Finally, to quantify individual differences in decision consistency during the Transfer phase, we estimated a softmax inverse temperature parameter *β* for each participant. Because no feedback was provided during Transfer, reward values for each symbol were fixed at their trained probabilities (A = 0.80, B = 0.20, C = 0.70, D = 0.30, E = 0.60, F = 0.40). For every choice trial, we computed the likelihood of selecting the observed option based on a standard softmax rule:

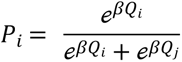

where *Q_i_*and *Q_i_*are the reward values of the chosen and unchosen stimuli, respectively. The total log-likelihood across all trials was defined as:

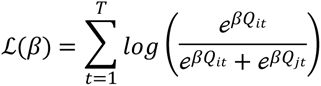

The value of *β* was estimated individually for each participant by maximizing this log-likelihood function using Brent’s method over the interval [0.001, 20]. Larger *β* values reflect more deterministic (exploitative) choices, whereas smaller values indicate more stochastic (exploratory) behavior.

These β estimates were compared between groups (Control vs. Gamer) using a linear regression.

## Results

### Sample Characteristics

Table 1 summarizes the means and standard deviations for weekly gaming hours, the Barratt Impulsiveness Scale (BIS-11), and the Sensitivity to Punishment and Sensitivity to Reward Questionnaire (SPSRQ) providing a descriptive overview of the sample characteristics. No group differences were observed in any measure, except for weekly gaming hours in the last six months, as expected.

**Table 1.** Sample Characterization.

| Variable | Control *M* (*SD*) | Gamer *M* (*SD*) | *t* | *p* | Cohen's *d* |
| --- | --- | --- | --- | --- | --- |
| **Weekly Gaming Hours** | 1.83 (1.87) | 20.56 (11.05) | -9.88 | **<.001\*\*** | -2.32 |
| **BIS-11** | 62.47 (6.66) | 59.94 (11.73) | 1.11 | 0.27 | 0.26 |
| **SPSRQ Reward** | 11.50 (5.17) | 10.61 (4.34) | 1.40 | 0.16 | 0.32 |
| **SPSRQ Punishment** | 12.81 (5.26) | 12.77 (4.82) | 0.02 | 0.98 | 0.005 |

*Note: BIS-11 = Barratt Impulsiveness Scale; SPSRQ= The Sensitivity to Punishment and Sensitivity to Reward Questionnaire. Significant comparisons are presented in bold \* $p \leq 0.05$ , \*\* $p < .001$*

### Learning Phase

The number of learning-test cycles required to reach the transfer criterion did not differ significantly between the Control and Gamer groups (*t*(67.50) = 1.84, *p* = .07). Although not statistically significant, controls completed slightly more learning-test cycles on average (*M* = 3.81) compared to Gamers (*M* = 3.17).

The generalized linear mixed-effects model (Figure 2, Table 2,) examining accuracy across the Learning phase revealed significant main effects of Block (*z* = 5.66, *p* < .001), indicating overall improved performance over time, and Pair, specifically highlighting lower accuracy for pair E/F compared to A/B (*z* = -4.38, *p* < .001). Crucially, there was a significant three-way interaction among Group, Block, and Pair, with gamers showing differential learning trajectories compared to controls, particularly for pairs C/D (*z* = -2.03, *p* = .043) and E/F (*z* = -2.66, *p* = .008). This suggests that while both groups improved over blocks, Gamers exhibited distinct learning patterns depending on stimulus difficulty.

**Figure 2.**
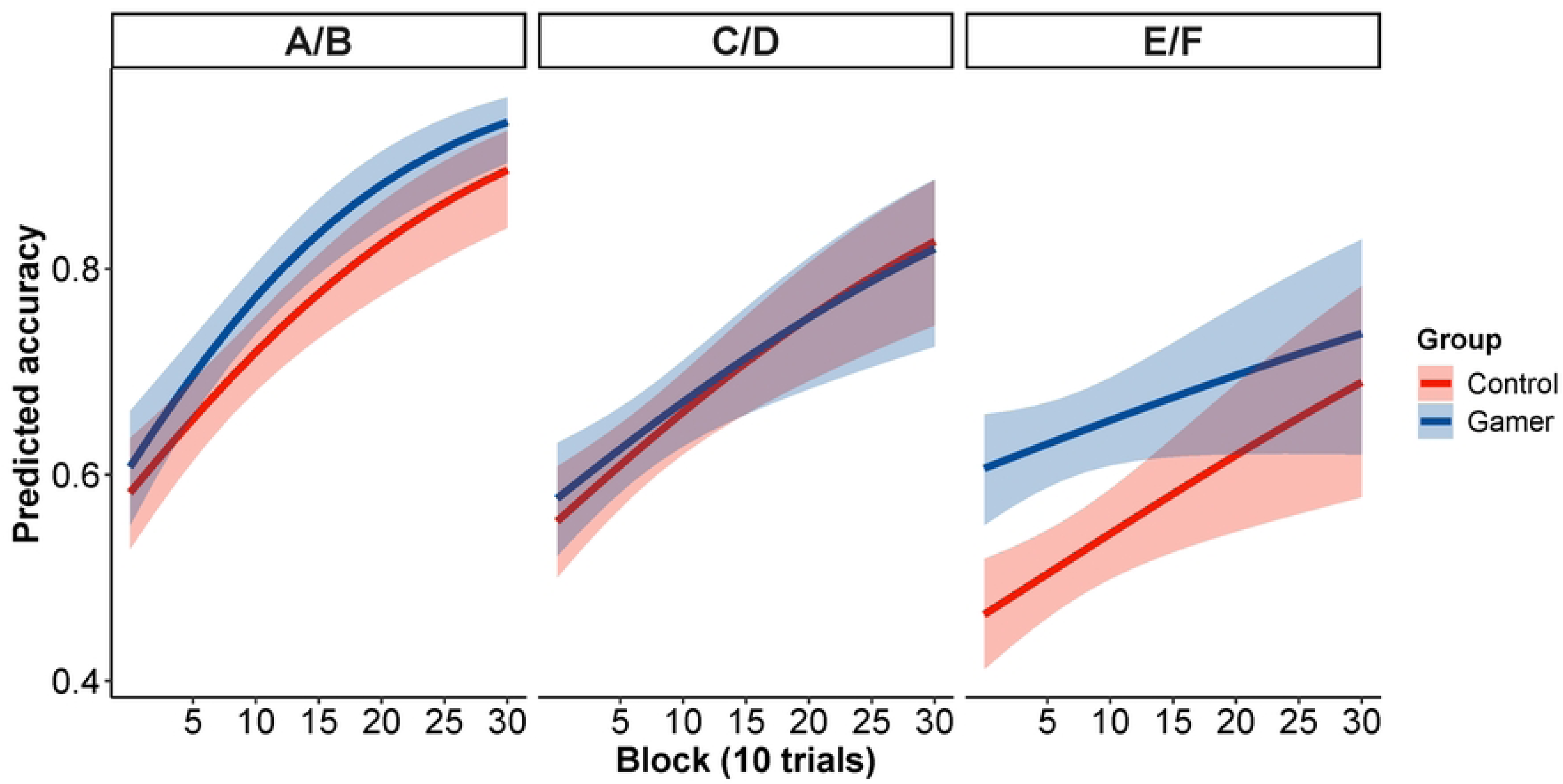
Learning curves. Predicted accuracy across training blocks (each block = 10 trials) for three stimulus conditions (A/B, C/D, and E/F), separated by experimental group. Lines represent model-based predicted accuracy for the Control (red) and Gamer (blue) groups across training. Shaded regions denote 95% confidence intervals.

**Table 2.**
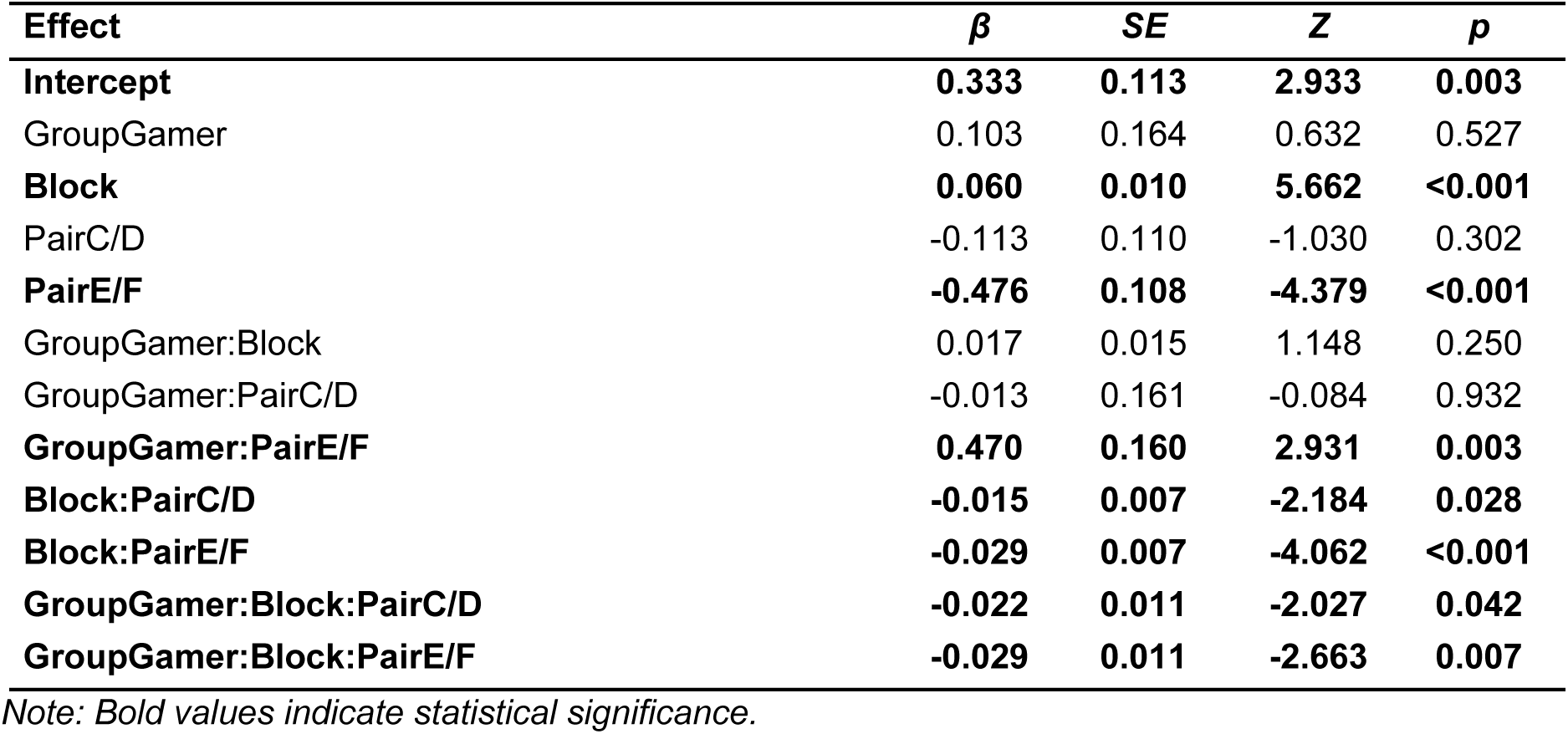
Model for accuracy across learning blocks.

| Effect | $\beta$ | SE | Z | p |
| --- | --- | --- | --- | --- |
| Intercept | **0.333** | **0.113** | **2.933** | **0.003** |
| GroupGamer | 0.103 | 0.164 | 0.632 | 0.527 |
| **Block** | **0.060** | **0.010** | **5.662** | **<0.001** |
| PairC/D | -0.113 | 0.110 | -1.030 | 0.302 |
| **PairE/F** | **-0.476** | **0.108** | **-4.379** | **<0.001** |
| GroupGamer:Block | 0.017 | 0.015 | 1.148 | 0.250 |
| GroupGamer:PairC/D | -0.013 | 0.161 | -0.084 | 0.932 |
| **GroupGamer:PairE/F** | **0.470** | **0.160** | **2.931** | **0.003** |
| **Block:PairC/D** | **-0.015** | **0.007** | **-2.184** | **0.028** |
| **Block:PairE/F** | **-0.029** | **0.007** | **-4.062** | **<0.001** |
| **GroupGamer:Block:PairC/D** | **-0.022** | **0.011** | **-2.027** | **0.042** |
| **GroupGamer:Block:PairE/F** | **-0.029** | **0.011** | **-2.663** | **0.007** |

*Note: Bold values indicate statistical significance.*

The mixed-effects logistic regression examining Win-Stay/Lose-Shift behavior (Figure 3, Table 3) revealed significant effects for previous outcome (z = -10.85, p < .001), indicating lower probability of staying with a choice after losses compared to wins, and accumulative reward (z = 2.62, p = .009), suggesting higher decision persistence with increasing reward accumulation. Additionally, a significant interaction between Group and accumulated reward was observed (z = 3.07, p = .002), indicating that gamers exhibited greater sensitivity to accumulated rewards in their decision-making processes compared to controls. Pair effects were also significant, with pairs C/D (z = -3.98, p < .001) and E/F (z = -5.92, p < .001) associated with lower stay probabilities, reflecting differential difficulty or uncertainty associated with these pairs. In sum, the steeper increase in the Gamer group suggests a heightened reliance on reward history to guide decision-making, indicative of more adaptive or efficient reinforcement learning strategies.

**Figure 3.**
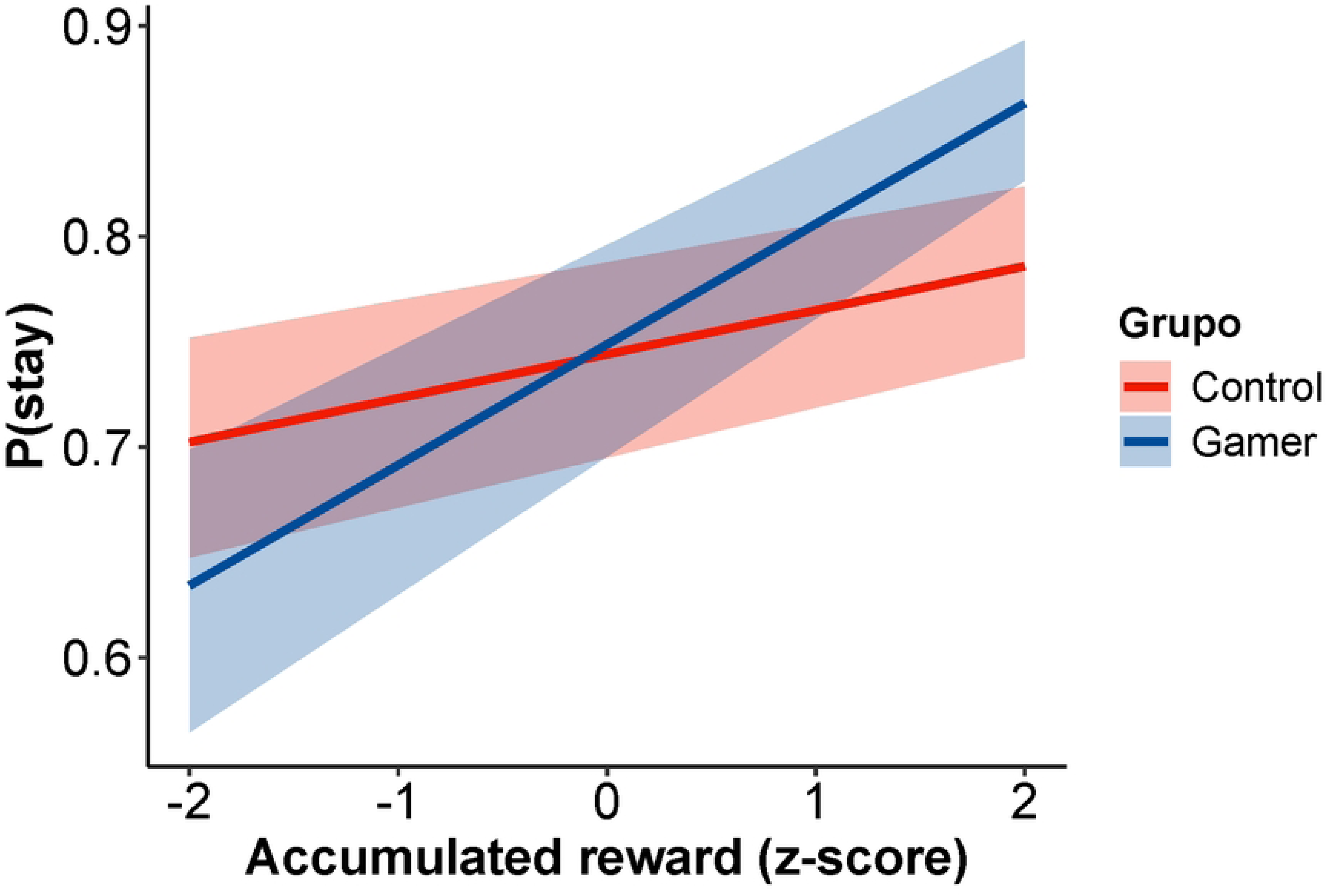
Win-Stay/Lose-Shift analysis. Predicted probability of staying with the same choice as a function of accumulated reward (z-scored), plotted separately for the Control (red) and Gamer (blue) groups. Shaded areas represent 95% confidence intervals.

**Table 3.** Win-Stay/Lose-Shift model.

| Effect | $\beta$ | SE | Z | p |
| --- | --- | --- | --- | --- |
| (Intercept) | **1.079** | **0.092** | **11.621** | **<0.001** |
| GroupGamer | 0.117 | 0.128 | 0.909 | 0.363 |
| **PreviousOutcomeLoss** | **-0.876** | **0.08** | **-10.851** | **<0.001** |
| **AccumulateReward** | **0.11** | **0.042** | **2.622** | **0.008** |
| **PairC/D** | **-0.175** | **0.044** | **-3.983** | **<0.001** |
| **PairE/F** | **-0.259** | **0.043** | **-5.918** | **<0.001** |
| GroupGamer:PreviousOutcomeLoss | 0 | 0.117 | -0.004 | 0.996 |
| **GroupGamer:AccumulateReward** | **0.212** | **0.069** | **3.069** | **0.002** |
| PreviousOutcomeLoss:AccumulateReward | -0.103 | 0.056 | -1.823 | 0.068 |
| GroupGamer:PreviousOutcomeLoss:AccumulateReward | -0.026 | 0.089 | -0.293 | 0.769 |

*Note: Bold values indicate statistical significance.*

The logistic mixed-effects regression analysis examining exploration behavior (Figure 4) showed significant main effects of Group (z = -2.19, p = .028), with gamers displaying a lower overall likelihood of exploring compared to controls, and global trial (z = -7.68, p < .001), indicating a general reduction in exploration across trials. The interaction between Group and global trial was not significant (z = 0.34, p = .736), suggesting a similar rate of exploration reduction over time for both groups. Consequently, although both groups exhibited a decrease in exploration over time, consistent with learning and strategy stabilization, the Gamer group demonstrated a consistently lower probability of exploration across the task, suggesting a more exploitative choice pattern compared to controls.

**Figure 4.**
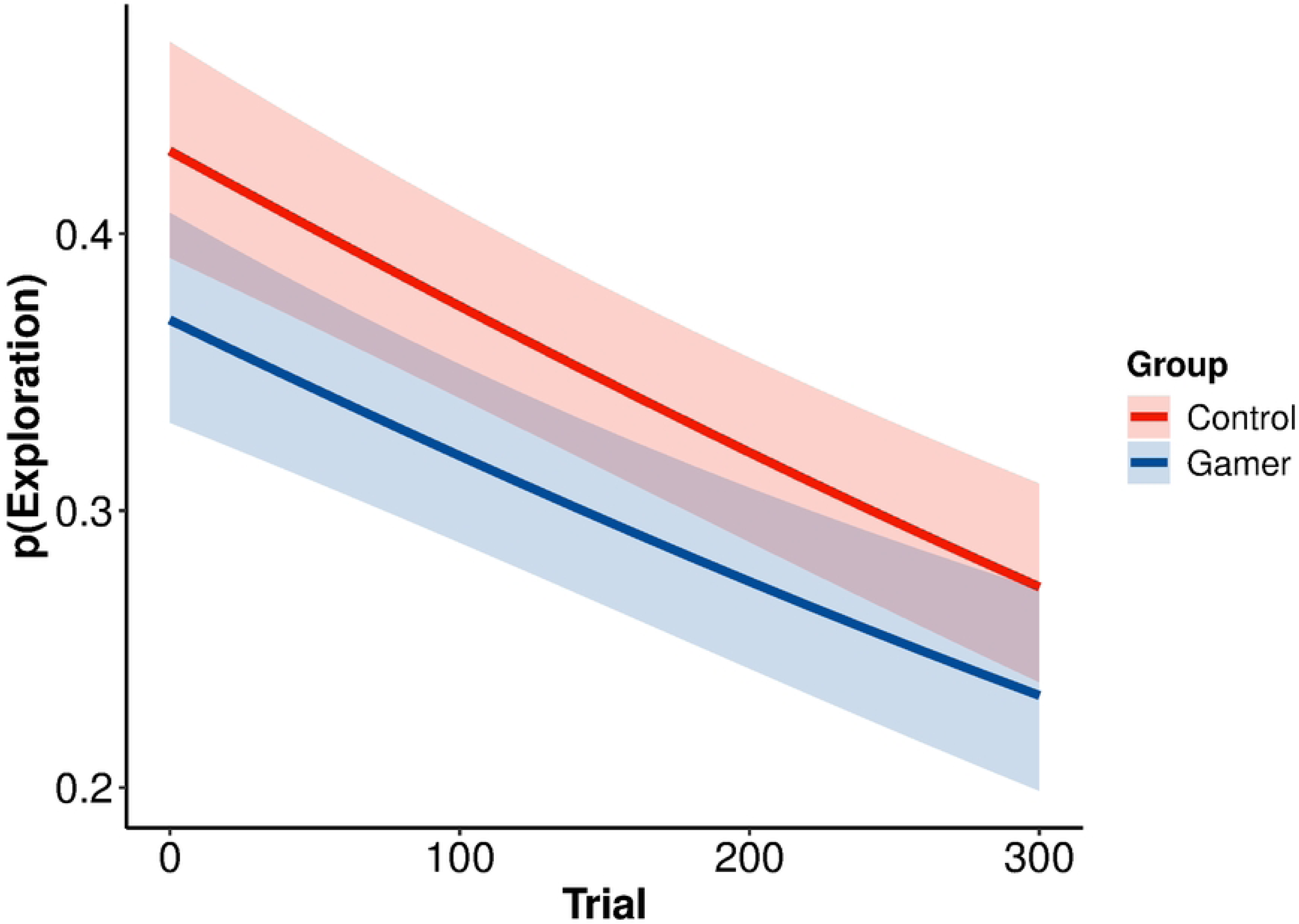
Exploration/Explotation analysis. Predicted probability of exploratory choices across trials for the Control (red) and Gamer (blue) groups. Shaded regions represent 95% confidence intervals.

### Test phase

The mixed-effects logistic regression analysis of accuracy during the Test phase (Figure 5) revealed significant main effects for Group (z = 1.99, p = .047), indicating that Gamers had higher overall accuracy than Controls, and Pair, with pair C/D (z = -4.47, p < .001) and E/F (z = -12.58, p < .001) showing significantly lower accuracy compared to pair A/B. The interaction between Group and Pair was significant for pair E/F (z = 2.80, p = .005), but not for the C/D pair (z = 1.09, p = .27), suggesting Gamers performed better specifically on the most challenging stimulus pair (E/F) compared to Controls.

**Figure 5.**
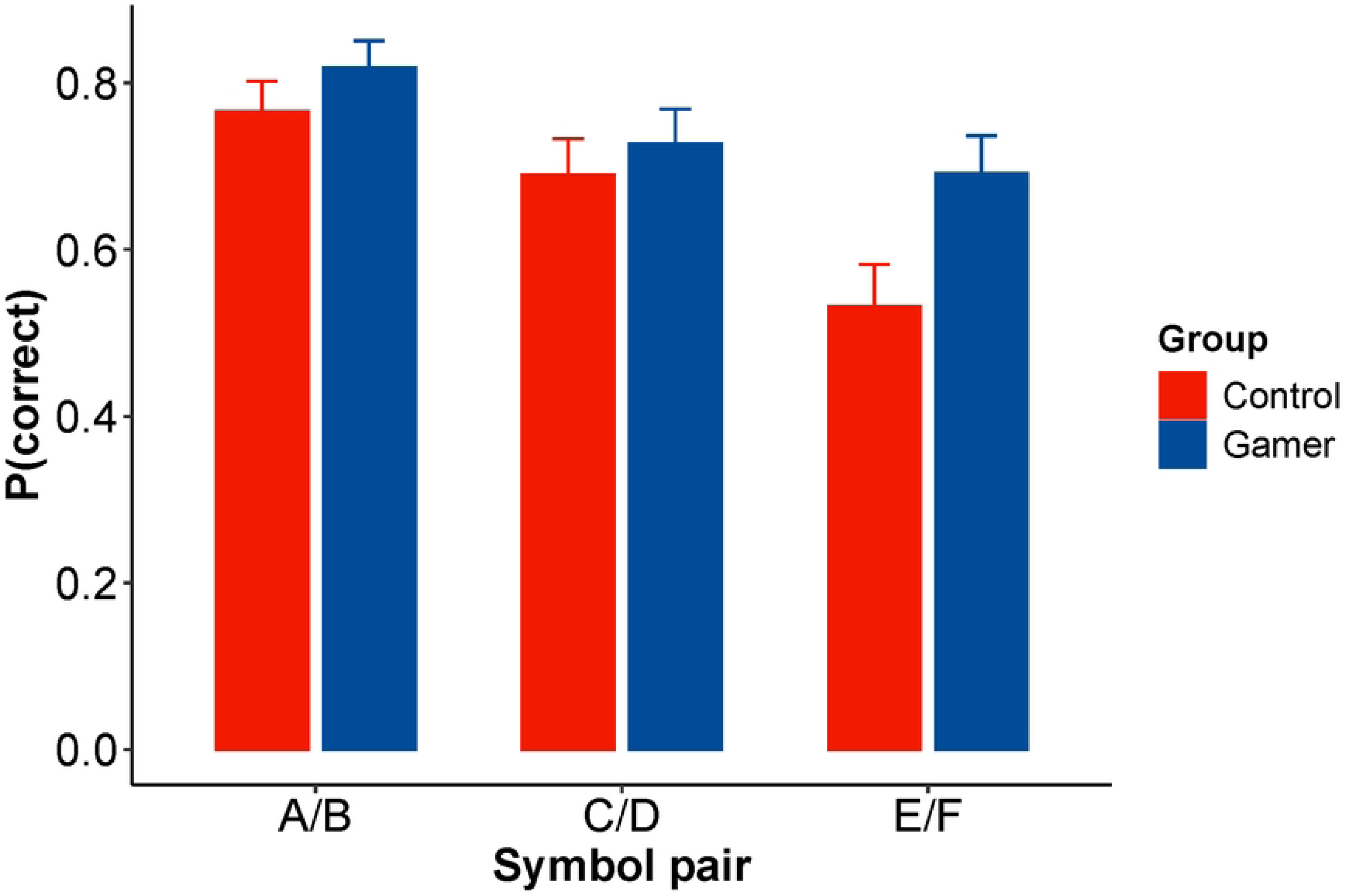
Correct responses on the Test phase. Mean probability of correct choices across symbol pairs (A/B, C/D, E/F) for Control (red) and Gamer (blue) groups, with error bars representing standard errors.

The linear mixed-effects model examining entropy during the Test phase revealed significant main effects for Pair, with higher entropy observed for pairs C/D (t = 2.39, p = .01) and E/F (t = 4.02, p < .001) compared to pair A/B. This indicates greater decisional uncertainty or variability for the more difficult stimulus pairs. There were no significant effects of Group (t = 0.95, p = .34) or interactions between Group and Pair (C/D: t = 1.03, p = .27; E/F: t = 1.12, p = .26), suggesting that decisional consistency did not differ substantially between Control and Gamer groups.

### Transfer phase

In the Transfer phase, separate logistic mixed-effects regressions for selecting stimulus A (choose_A) and avoiding stimulus B (avoid_B) showed no significant main effect of Group (choose_A: z = 0.97, p = .33; avoid_B: z = 1.50, p = .13), suggesting comparable performance between groups in transferring learned stimulus values. Furthermore, the Valence Learning Bias Index (VLBI), measuring preference toward approach versus avoidance behavior, did not differ significantly between groups (t(70) = -0.26, p = .798).

Finally, the inverse temperature parameter (β), which quantifies choice consistency or exploitation during the Transfer phase, was analyzed with a linear regression model. No significant differences between Control and Gamer groups were observed (t(70) = 1.19, p = .23), indicating that both groups exhibited comparable levels of choice consistency.

## Discussion

The present study investigated how habitual video game play relates to reinforcement learning dynamics, emphasizing trial-by-trial adaptation across probabilistic contingencies, followed by consolidation and transfer without feedback.

As expected, participants in the gamer group reported significantly more weekly gaming hours than those in the Control group, with a very large effect size, confirming the robustness of the group classification. In contrast, no significant differences were found between groups in impulsivity levels (BIS-11) or in sensitivity to reward and punishment (SPSRQ), variables that have been linked to reinforcement learning performance [18,19]. These findings suggest that the groups were comparable in their general self-perceived sensitivity to reward and punishment, as well as in self-regulatory traits, and that observed differences in task performance are unlikely to be explained by dispositional impulsivity or sensitivity to general reinforcement cues.

Based on previous findings that gaming experience enhances probabilistic inference [10] and probabilistic categorization learning outcomes [8], we aimed to disentangle, through trial-by-trial modeling, how individuals update value expectations based on feedback.

Consistent with our hypothesis, during the learning phase gamers demonstrated superior performance in these dynamics. Although the number of learning–test cycles required to reach criterion did not significantly differ between groups, gamers completed slightly fewer cycles on average, suggesting a trend toward more efficient learning. A significant three-way interaction between Group, Block, and Pair revealed that gamers displayed distinct learning trajectories, particularly for the more difficult discrimination pairs (C/D and E/F). This pattern reflects an enhanced capacity to integrate probabilistic feedback and adjust responses under increasing task demands, consistent with Schenk’s finding [8] that gamers performed better in learning categorization tasks under conditions of higher uncertainty.

Our trial-level analyses in the learning phase provided a more detailed view of how gamers modulated behavior over time. The significant Group × Reward interaction in the Win– Stay/Lose–Shift analysis indicated that gamers were more likely to persist in rewarded choices as the accumulated reward increased. This behavioral pattern is consistent with reinforcement learning frameworks in which action preferences are updated incrementally based on the difference between the obtained reward and a reference level, such as the running average reward. When this difference is positive, the probability of repeating the same choice increases—a process formalized in gradient bandit methods and temporal-difference learning, where value estimates are adjusted trial-by-trial toward the experienced outcome [1]. Neurobiological accounts parallel these principles, with converging evidence that phasic midbrain dopamine activity encodes temporal-difference–like reward-prediction errors: brief increases in firing when outcomes are better than expected, no change when outcomes match expectations, and transient decreases when expected rewards are omitted. These signals act as teaching signals for synaptic modification and bias subsequent action selection toward repeating rewarded choices, providing a plausible mechanism by which positive feedback shapes ongoing decision policies [20,21].

Thus, these learning advantages may be supported by frequent engagement of the mesolimbic-cortical dopaminergic system during video game play, which has been shown to play a central role in encoding reward prediction errors and refining value representations [11,12].The more stable and less exploratory behavior patterns observed in gamers particularly for the hardest pair (E/F), reflect efficient exploitation of learned reward contingencies rather than behavioral rigidity. This is consistent with prior findings that gamers outperform non-gamers in cognitive flexibility tasks like the Wisconsin Card Sorting Test [22] and in decision-making under uncertainty, as shown in the Iowa Gambling Task [22,23].

The exploration–exploitation analysis further supported this view, revealing that gamers engaged in consistently less exploratory behavior across trials. While both groups reduced exploration over time, gamers’ overall lower exploration rates suggest earlier stabilization of decision policies and a more exploitative reinforcement learning strategy. These behavioral patterns are likely supported not only by neural adaptations, but also by the cognitive demands embedded in gaming environments [12]. Video games usually require rapid evidence accumulation, outcome monitoring, and adaptive strategy selection in response to noisy feedback. These parallels suggest that video games may operate as a naturalistic reinforcement learning context, shaping both neural and cognitive systems that support feedback integration and decision-making under uncertainty.

In the test phase, where participants had to apply previously learned contingencies without receiving feedback, gamers outperformed controls in overall accuracy, particularly on the most difficult pair (E/F). A significant Group × Pair interaction indicated that gamers retained and applied learned associations more effectively under high ambiguity. This result aligns with Schenk et al.’s [8] WPT findings and, importantly, extends them: our data suggest that gaming experience may facilitate the consolidation of learned value representations, supporting performance even when reinforcement (feedback) is withheld. On the other hand, Entropy analyses revealed greater decision uncertainty for the more difficult stimulus pairs in both groups, but no group differences emerged. This finding suggests that although gamers were more accurate, they did not exhibit greater decisional consistency. Their advantage appears to stem from more precise internal value representations, rather than from reduced variability in their choices

Such reinforcement-related advantages also have potential relevance for learning and adaptation in everyday contexts. Educational settings often rely on incremental skill development supported by corrective feedback. Incorporating game-like elements, such as immediate reinforcement, adaptive difficulty, and variable reward schedules may enhance student engagement and improve feedback integration. Evidence from educational psychology supports this view: intrinsically integrated educational games have been shown to promote both learning and motivation by aligning task goals with gameplay mechanics [24]. Similarly, a meta-analysis by Wouters et al. [25] revealed that serious games yield superior outcomes compared to conventional instruction, especially when they include mechanisms like real-time feedback and adaptive challenge. These findings suggest that game-based dynamics may amplify the benefits of reinforcement-driven learning by providing a structured yet flexible framework for performance monitoring and strategic adjustment. While Bembenutty and Karabenick [16] emphasized the role of feedback and self-regulation in academic success, these more recent studies illustrate how gamified feedback structures can further optimize cognitive engagement in educational environments. Building on this, our study contributes by examining whether habitual video game experience is associated with measurable differences in the reinforcement learning processes that could support such transfer, providing insight into the mechanisms that sustain these advantages.

In the transfer phase, no significant differences emerged between groups in selecting the most rewarded stimulus or avoiding the most punished one. The Valence Learning Bias Index (VLBI) did not differ between gamers and controls, nor did the inverse temperature parameter (β), indicating similar levels of choice consistency and value generalization across novel stimulus recombinations [26,27].

These findings indicate that the reinforcement learning advantages observed in gamers— such as heightened reward sensitivity, improved performance under uncertainty, and earlier policy stabilization—may be most evident in dynamic, feedback-rich contexts [8,28]. In contrast, when feedback is removed and learning must be generalized to novel combinations, both groups appear to rely on similar motivational and decisional processes. This dissociation suggests that gaming influences flexible, state-dependent reinforcement strategies more than stable, trait-like learning biases.

Interestingly, these differences in reinforcement learning trajectories may also have implications for individual variability in adaptation across domains. For example, reinforcement profiles have been linked to clinical traits, with punishment sensitivity associated with anxiety and compulsivity, and reward sensitivity with impulsivity and diminished self-regulation [13,14]. While our results did not reveal group differences in valence bias, the enhanced reward-driven persistence observed in gamers raises important questions about how repeated exposure to reward-rich environments might shape the calibration of motivational systems. Depending on the context, such tuning may promote more efficient learning—as suggested by longitudinal findings of gaming training-related increases in reward responsiveness [29]—or, alternatively, contribute to maladaptive outcomes, as indicated by studies on Internet Gaming Disorder showing heightened sensitivity to reward cues and impaired regulatory control [30].

Taken together, the present findings highlight that habitual video game play is associated with more efficient reinforcement learning strategies during acquisition and testing, but these effects may not extend to motivational biases in generalized transfer contexts. Gaming appears to promote flexible value updating and feedback integration under uncertainty— capacities that may hold relevance for both educational design and the understanding of individual variability in adaptive learning.

### Considerations

Several considerations should be noted. A plausible explanation for the absence of a bias in gamers during the transfer phase is that the block configuration always paired the most rewarded option (A) and the most punished option (B) with other alternatives. Although gamers outperformed controls in the learning and test phases with greater reward sensitivity, especially under high-uncertainty conditions, controls also discriminated A and B well. As a result, the current structure may have masked any bias in these high-certainty contrasts.

Future research could address this by designing transfer blocks that include lower-certainty symbol combinations, where gamers might display the hypothesized bias.

Larger samples would also clarify near-significant trends (e.g., learning-test cycles) and allow exploration of moderating factors such as gaming genre or cumulative playtime. Future studies should examine whether reinforcement learning dynamics and biases differ across video game genres. Specifically, action gamers demonstrate improved probabilistic inference, allowing them to make more accurate decisions based on uncertain or incomplete information [31]. In addition, recent findings indicate that such players exhibit superior “learning to learn” abilities, characterized by a heightened capacity to detect changes in environmental regularities and adjust their learning strategies accordingly [32]. Strategy games, such as real-time strategy (RTS) games, emphasize planning, multitasking, and dynamic decision-making. These features have been shown to support enhancements in cognitive flexibility, a core aspect of cognitive control [33]. Similarly, expertise in multiplayer online battle arena (MOBA) games has been associated with superior inhibitory control and greater neural efficiency in frontoparietal networks involved in executive functioning [34].

Because our gamer sample included participants from multiple genres (see S1 Table), the observed effects likely reflect reinforcement-relevant design features common across commercial video games—adaptive difficulty that responds to performance, instant and continually provided feedback that supports self-monitoring and rapid policy adjustment, and goal-directed, challenge-rich environments that deliver contingent rewards—rather than mechanics unique to a single genre [35]. Nonetheless, genre-stratified analyses in future work could test whether these learning benefits are genre-dependent or generalize across contexts. Controlled, genre-based training interventions would further assess causality and specificity.

Moreover, most research describes how video games sustain continuous, goal-contingent rewards. Yet many games also engage avoidance learning by imposing strong penalties for failure. Although some experiments have tested how such penalties influence player experience, learning, and interest, their primary focus has been game design rather than cognitive mechanisms [36,37]. This underexplored feature could contribute to learning biases on the Probabilistic Selection Task (PST). To determine its impact, longitudinal training studies that systematically manipulate failure-penalty severity in specific game contexts are needed.

Finally, neuroimaging and computational modeling approaches could further elucidate how experience-driven plasticity in dopaminergic and fronto-striatal circuits underpins these behavioral effects. Moreover, intervention studies incorporating controlled gaming paradigms could determine whether structured exposure to game-like reinforcement schedules causally enhances feedback-based learning efficiency.

## Conclusion

Our findings indicate that habitual video game play is associated with enhanced trial-by-trial reinforcement integration, greater reward sensitivity in the learning dynamics, and superior performance in conditions of high ambiguity during acquisition and testing phases of the PST. However, these advantages did not extend to measures of learning biases toward reward or punishment in the transfer phase, suggesting that gaming primarily sharpens dynamic reinforcement-driven learning processes. By showing how video games, through their reliance on continuous feedback, probabilistic contingencies, and adaptive decision-making engage cognitive mechanisms central to reinforcement learning theory, this study supports their utility as a model for investigating how feedback-rich environments shape learning. These insights not only advance our understanding of reinforcement learning in naturalistic contexts but also highlight potential educational and clinical applications of feedback-driven training approaches modeled on gaming environments.

## Supporting Information

Table S1 S2. File

## Acknowledgments

The author would like to thanks all participants for their time and commitment to this study.

## Author Contributions

**Conceptualization:** LALA AQAC AJQC

**Data Curation:** LALA AQAC AJQC AAO

**Methodology:** LALA AQAC AJQC AAO

**Project Administration:** LALA

**Resources:** LALA SLQB

**Formal analysis:** AQAC

**Investigation:** AJQC AAO

**Validation:** AQAC SLQB ALJP

**Visualization:** LALA AQAC SLQB ALJP

**Supervision:** LALA SLQB ALJP

**Software:** AQAC

**Writing ± original draft:** LALA AQAC

**Writing ± review & editing:** LALA AQAC SLQB ALJP

